# Synergistic and offset effects of fungal species combinations on plant performance

**DOI:** 10.1101/2021.05.09.443080

**Authors:** Yoshie Hori, Hiroaki Fujita, Kei Hiruma, Kazuhiko Narisawa, Hirokazu Toju

## Abstract

In natural and agricultural ecosystems, survival and growth of plants depend substantially on microbes in the endosphere and rhizosphere. Although numerous studies have reported the presence of plant-growth promoting bacteria and fungi in below-ground biomes, it remains a major challenge to understand how sets of microbial species positively or negatively affect plants’ performance. By conducting a series of single- and dual-inoculation experiments of 13 endophytic and soil fungi targeting a Brassicaceae plant species, we here evaluated how microbial effects on plants depend on presence/absence of co-occurring microbes. The comparison of single- and dual-inoculation experiments showed that combinations of the fungal isolates with the highest plant-growth promoting effects in single inoculations did not yield highly positive impacts on plant performance traits (e.g., shoot dry weight). In contrast, pairs of fungi including small/moderate contributions to plants in single-inoculation contexts showed the greatest effects on plants among the 78 fungal pairs examined. These results on the offset and synergistic effects of pairs of microbes suggest that inoculation experiments of single microbial species/isolates can result in the overestimation or underestimation of microbial functions in multi-species contexts. Because keeping single-microbe systems in outdoor conditions is impractical, designing sets of microbes that can maximize performance of crop plants is an important step for the use of microbial functions in sustainable agriculture.

## INTRODUCTION

Plants in natural and agricultural ecosystems are associated with diverse taxonomic groups of microbes, forming both positive and negative interactions with the microbiomes (Lundberg et al., 2012; Peay et al., 2016; Busby et al., 2017; Toju et al., 2018b). In particular, bacteria and fungi found within and around root systems have been reported as key determinants of plants’ survival and growth (Hiruma et al., 2016, 2018; Castrillo et al., 2017; Trivedi et al., 2020). A number of rhizosphere bacteria, for example, are known to stimulate plants’ growth by producing phytohormones (Lugtenberg and Kamilova, 2009; Bhattacharyya and Jha, 2012; Finkel et al., 2020). Mycorrhizal fungi are ancient symbionts of land plants (Remy et al., 1994; Taylor et al., 1995), providing soil phosphorus and/or nitrogen to their hosts (Richardson et al., 2009; Tedersoo et al., 2010; Jansa et al., 2019). Moreover, a growing number of studies have shown that diverse clades of endophytic and soil fungi support host plants by provisioning inorganic/organic forms of nutrients (Usuki and Narisawa, 2007; Newsham, 2011; Hiruma et al., 2016), activating plant immune systems (van Wees et al., 2008; Pieterse et al., 2014), and suppressing populations of pathogens/pests in the rhizosphere (Narisawa et al., 2004; Khastini et al., 2012; Gu et al., 2020). Thus, developing scientific bases for maximizing the benefits from those plant-associated microbiomes is an essential step for fostering sustainable agriculture and restoring forest/grassland ecosystems (Bulgarelli et al., 2013; Carlström et al., 2019; Wagg et al., 2019; Saad et al., 2020).

One of the major challenges in utilizing plant-associated microbiome functions is to design sets of microbial species/isolates (Vorholt et al., 2017; Paredes et al., 2018; Toju et al., 2018a; Wei et al., 2019). While a single microbial species or isolate can have specific functions in promoting plant growth, broader ranges of positive effects on plants are potentially obtained by introducing multiple microbial species/isolates (Wang et al., 2011; Ważny et al., 2018; He et al., 2020). For example, a fungal species degrading organic nitrogen (Newsham, 2011) and that suppressing soil pathogens (Vinale et al., 2008) may provide plants with a broader spectrum of physiological functions than each of them alone, potentially having additive or synergistic effects on the growth of their hosts. Meanwhile, sets of microbes trying to colonize the plant endosphere or rhizosphere may compete for resources/space (Kennedy et al., 2009; Werner and Kiers, 2015; Toju et al., 2016) or inhibit each other’s growth (Helfrich et al., 2018), making their impacts on host plants more negative than that observed in single-inoculation conditions (i.e., offset effects) (Nelson et al., 2018).

Given that multiple microbial species inevitably interact with a single plant in agroecosystems (Toju et al., 2018a), knowledge of those synergistic and offset effects in plant-associated microbiomes is crucial for optimizing microbial functions in agriculture.

A starting point for designing sets of microbes is to use the information of single-inoculation assays, in each of which a single microbial species/isolate is introduced to a target plant species/variety (Ahmad et al., 2008; Harbort et al., 2020). Through this initial assay, respective species/isolates are scored in terms of their functions (e.g., plant-growth promotion effects) in single-inoculation conditions (Nara, 2006; Dai et al., 2008; Taurian et al., 2010; Tsolakidou et al., 2019). The next step is to consider how we can use these single-inoculation scores for designing sets of microbes that potentially promote plant growth in synergistic ways. As the number of combinations inflates with that of constituent species/isolates [e.g., {N × (N – 1)}/2 combinations in two-species systems], prioritizing candidate species/isolate combinations based on single-inoculation results is an important step (Paredes et al., 2018; Toju et al., 2018a, 2020). The simplest way of exploring best sets of microbes is to combine microbes with highest single-inoculation scores. This strategy of combining microbes in highest ranks is promising if synergistic (or additive) effects are common in plant-associated microbiomes. In contrast, if offset effects of multiple microbes on plant performance are ubiquitous, alternative strategies for exploring species/isolate combinations are required to maximize benefits from plant-associated microbiomes.

In this study, we tested the hypothesis that synergistic effects on plant growth are common in below-ground fungal biomes in a series of single- and dual-inoculation experiments. By using 13 endophytic/soil fungal species belonging to various taxonomic groups, we first evaluated their basic effects on plant growth in a single-inoculation experiments with a Brassicaceae species (*Brassica rapa* var. *perviridis*). We also performed dual-inoculation experiments for all the 78 possible combinations of the fungal species and then evaluated the performance of the combinations in light of single-inoculation results. The data then provided a platform for testing whether plant-growth promoting effects exceeding those of all the single-inoculation conditions are attainable in dual-inoculation conditions. We further examined whether such synergistic effects could be obtained with “high ranker × high ranker” combinations or in other types of combinations. Overall, this study provides a basis for understanding to what extent plant-growth promotion effects of microbiomes can be expected from the information of single-species inoculations, illuminating the potential importance of “non-additivity” in multi-microbe contexts.

## MATERIALS AND METHODS

### Fungal isolates for inoculation experiments

In the inoculation experiments detailed below, we used diverse fungal species isolated from plant roots or soil (Table 1). Among the 13 fungal isolates used (Table 1; Supplementary Data S1), some are reported as endophytic fungi promoting host plant growth [e.g., *Colletotrichum tofieldiae, Cladophialophora chaetospira*, and *Veronaeopsis simplex*] in previous studies (Usuki and Narisawa, 2007; Hiruma et al., 2016; Guo et al., 2018). In addition, a species of *Trichoderma* with growth-promotion effects on tomato (*Solanum lycopersicum*) and *Brassica* plants (Toju et al., 2020) was used in the experiment. To gain insights from a broad ecological spectrum of fungi in the experiments, isolates belonging to diverse genera were selected from the ca. 3,500 fungal isolates maintained in the culture collection of Centre for Ecological Research, Kyoto University. Putative functional groups of these fungi were inferred using the FUNGuild program (Nguyen et al., 2016) as shown in Table 1. Note that such profiling information based on ecological guild databases should be interpreted with caution: even in a fungal genus embracing a number of plant pathogenic species, some species can have positive impacts on plants (Radhakrishnan et al., 2015; Hiruma et al., 2016).

**TABLE 1.** Fungal isolates used in the inoculation experiments.

| Isolate | Abbreviation | Phylum | Class | Order | Family | Genus | Guild | Blast top-hit | E value | Per. Ident | Accession |
| --- | --- | --- | --- | --- | --- | --- | --- | --- | --- | --- | --- |
| *Phoma* sp. KUCER00000052 | pho_0052 | Ascomycota | Dothideomycetes | Pleosporales | Didymellaceae | *Phoma* | P ES | *Phoma leveillei* | 9.00E-123 | 99.6% | KY827373.1 |
| *Alternaria* sp. KUCER00001239 | alt_1239 | Ascomycota | Dothideomycetes | Pleosporales | Periconiaceae | *Alternaria* | P ES | *Alternaria broccoli-italicae* | 2.00E-123 | 100.0% | MH374617.1 |
| *Curvularia* sp. KUCER00000077 | cur_0077 | Ascomycota | Dothideomycetes | Pleosporales | Periconiaceae | *Curvularia* | P | *Curvularia coatesiae* | 4.00E-126 | 100.0% | MK804384.1 |
| *Setosphaeria* sp. KUCER00000031 | set_0031 | Ascomycota | Dothideomycetes | Pleosporales | Periconiaceae | *Setosphaeria* | P E | *Setosphaeria pedicellata* | 1.00E-126 | 100.0% | LT837452.1 |
| *Stemphylium* sp. KUCER00000804 | ste_0804 | Ascomycota | Dothideomycetes | Pleosporales | Periconiaceae | *Stemphylium* | P S | *Stemphylium lycopersici* | 2.00E-125 | 100.0% | MN386223.1 |
| *Veronaeopsis simplex* Y34 | ver_0232 | Ascomycota | Dothideomycetes | Venturiales | Sympoventuriaceae | *Veronaeopsis* | E | *Veronaeopsis simplex* | 5.00E-125 | 100.0% | MH865233.1 |
| *Cladophialophora chaetospora* M4006 | cla_0230 | Ascomycota | Eurotiomycetes | Chaetothyriales | Herpotrichiellaceae | *Cladophialophora* | E | *Cladophialophora chaetospora* | 3.00E-123 | 99.6% | LC077702.1 |
| *Aspergillus* sp. KUCER00000917 | asp_0917 | Ascomycota | Eurotiomycetes | Eurotiales | Aspergillaceae | *Aspergillus* | S | *Aspergillus terreus* | 7.00E-124 | 99.6% | MH124236.1 |
| *Colletotrichum tofieldiae* MAFF 712334 | col_0223 | Ascomycota | Sordariomycetes | Glomerellales | Glomerellaceae | *Colletotrichum* | P E | *Colletotrichum tofieldiae* | 2.00E-125 | 100.0% | KX069824.1 |
| *Trichoderma* sp. KUCER00000218 | tri_0218 | Ascomycota | Sordariomycetes | Hypocreales | Hypocreaceae | *Trichoderma* | PFES | *Trichoderma asperellum* | 5.00E-125 | 100.0% | MT530021.1 |
| *Fusarium* sp. KUCER00000983 | fus_0983 | Ascomycota | Sordariomycetes | Hypocreales | Nectriaceae | *Fusarium* | P ES | *Fusarium oxysporum* | 5.00E-125 | 100.0% | MT610995.1 |
| *Tolypocladium* sp. KUCER00000289 | tol_0289 | Ascomycota | Sordariomycetes | Hypocreales | Ophiocordycipitaceae | *Tolypocladium* | FE | *Tolypocladium album* | 9.00E-123 | 99.6% | LC386577.1 |
| *Mucor* sp. KUCER00000113 | muc_0113 | Mucoromycota | - | Mucorales | Mucoraceae | *Mucor* | S | *Mucor abundans* | 1.00E-125 | 100.0% | MK164195.1 |

For each fungal isolate, taxonomy, functional guild information inferred by the FUNGuild database (P, plant pathogen; F, fungal pathogen; E, endophyte; S, saprophyte), and NCBI BLAST top-hit results of the ITS sequences are indicated for each isolate.

### Fungal inocula

Prior to the inoculation experiments, fungal inocula were prepared. Each of the 1.3-L high-density polyethylene bags with air-conditioning filters (Shinkoen Co. Ltd., Mino-kamo) was filled with the mixture of 60-cm^3^ wheat bran (Tamagoya Shoten), 60-cm^3^ rice bran, 180-cm^3^ leaf mold (Akagi Gardening Co., Ltd., S1), and 70-mL distilled water. The filled culture bags were sealed with a heat sealer (ANT-300, AS ONE Corporation, Osaka) and they were autoclaved three times at 121 °C for 30 min with 24 h intervals. For each fungal isolate, approximately ten pieces of mycelial disks (8.0 mm in diameter) was then transferred from 1/2 CMMY medium (cornmeal agar, 8.5 g/L; malt extract, 10.0 g/L; yeast extract, 1.0 g/L) (Becton, Dickinson and Co.) to the autoclaved substrate and the fungal culture bag was incubated at room temperature (approximately 25 °C) for 10–21 days until it was filled with mycelia. In addition to the 13 fungal inocula, a mock inoculum without fungi was prepared as a control.

Each of the fungal/control inocula was mixed with autoclaved potting soil consisting mainly of fermented bark, peat moss, and coconut peat [“Gin-no-tsuchi”; Total N, 0.41 % (w/w); P2O5, 0.62 %; K2O, 0.34 %; Kanea Inc., Takamatsu] by the proportion of 1:9. The mixed soil was then transferred into plastic cell trays: the size of each cell in the trays was 49 mm × 49 mm × 56.5 mm. Plant seeds were then introduced into the cell trays as detailed below.

### Inoculation experiments

The “Komatsuna Wase” variety of *Brassica rapa* var. *perviridis* (Atariya Noen Co. Ltd., Katori) was used as the target plant in the inoculation experiments. Before inoculation, the seeds of *Brassica* were surface sterilized by being shaken in 70 % ethanol solution for 1 min and then in 1 % sodium hypochlorite solution for 1.5 min. The seeds were then rinsed three times in distilled water. They were subsequently placed on 1 % agar petri dishes and incubated at 23 °C in the dark for 24-26 h until rooting. The rooted seeds were transferred to the inoculum-mixed soil on the following day: two seeds were introduced into each of the 20 or more replicate cells for each single inoculation experiment. The cell trays were maintained in the laboratory with the 16hL/8hD light condition at 25 °C. The plants were watered 3-4 times a week. The locations of the cell trays were rotated to equalize plants’ growing conditions.

In addition to the above single-inoculation experiments, dual-inoculation experiments were performed for all the 78 possible combinations of the 13 fungal isolates. For each pair of fungal isolate, their inocula were mixed by the proportion of 1 : 1, collectively constituting 1/10 volume of the total soil volume within the cell pots. Two *Brassica* seeds were then introduced into each of the 20 replicate cell pots and they were kept in the laboratory conditions detailed above. Due to the large number of treatments and replicates as well as the limited spatial capacity of the laboratory, the inoculation experiments were split into several experimental rounds (up to 13 single/dual/control treatments per round; see Supplementary Data S2 for the information of experimental rounds). To take into account potential difference of micro-environmental conditions among the experimental rounds, a control (mock inoculum) treatment was included in every round in order to standardize plant growth responses throughout the study (see below for the calculation of a standardized growth index).

After seven days, the ratio of geminating seeds to introduced seeds (i.e., germination rate) was recorded for each single/dual/control treatment. The seedlings were randomly thinned to one seedling per cell and they were kept in the same environmental conditions for another two weeks. The 21-day old *Brassica* plant samples were harvested to evaluate their responses to fungal inoculations. For all the replicate samples, shoot dry weight (above-ground biomass) and the number of mature leaves (> 20 mm in length) were recorded. For the measurement of shoot dry weight, plant samples were oven-dried at 60 °C for at least 72 h. Leaves longer than 20 mm were also subjected to SPAD measurements to infer chlorophyll concentrations using a SPAD-502Plus meter (Konica Minolta, Inc., Tokyo) (Netto et al., 2005; Zhu et al., 2012). For each of the randomly-selected 15 plant samples per treatment, the SPAD readings at three points were averaged. While shoot dry weight and the number of mature leaves are metrics of plant total biomass, SPAD readings are often regarded as (weak) indicators of foliar nitrogen concentrations (Chang and Robison, 2003; Esfahani et al., 2008).

### Plants’ growth responses

To standardize the variables representing plants’ responses to fungal inoculations, we proposed a standardized growth index as follows:

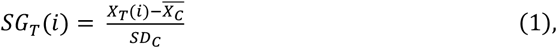

where *X*_*T*_(*i*) is a measurement of a target trait of a plant sample *i* in a target single-/dual-inoculation treatment, while 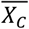 and *SD*_*C*_ are the mean and standard deviation of plant traits (variables) observed in the control samples of the focal experimental round, respectively. In terms of basic statistics assuming the Gaussian distribution, the standardized growth index [*SG*_*T*_(*i*)] values less than -1.96 and those larger than 1.96 roughly represented plant performance outside the 95 % confidence intervals of the control samples in the same experimental rounds, providing an intuitive criterion for comparing results within/across inoculation experiments (see Supplementary Figure S1 for relationship between the standardized growth index values and false discovery rates). This standardized growth index was calculated for each of the three plant variables representing plant performance: the number of mature leaves, shoot dry weight, and SPAD readings.

### Synergistic and offset effects

Based on the standardized growth index, we evaluated potential synergistic effects in dual inoculations of two fungal isolates in comparison to single-inoculation results. For a replicate plant sample inoculated with a pair of fungal isolates A and B, the index representing deviation from the maximum effects in single inoculations is calculated as follows:

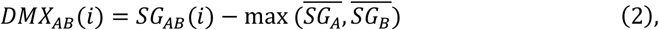

where *SG*_*AB*_(*i*) is the ce plant in the dual inoculation treatment, while 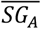 and 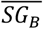 are means of standardized growth index values for the single inoculation of fungal isolates A and B, respectively. By definition, when there are synergistic effects of the presence of two fungal isolates [i.e., 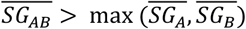], the mean of the deviation index over replicate plant samples 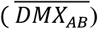 is larger than zero. Likewise, to evaluate offset effects in dual inoculations of two fungal isolates, an index representing deviation from the minimum effects in single inoculations was defined as follows:

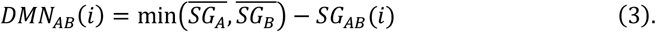

When there are offset effects [i.e., 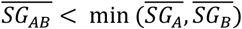] for a focal pair of fungi, mean of the offset effect index over replicate samples 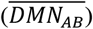 is larger than zero.

We further developed a simple index for evaluating deviations of observed dual-inoculation results from those expected as intermediate results of single inoculations. For a pair of fungal isolates A and B, the index for deviation from intermediate effects is calculated as follows:

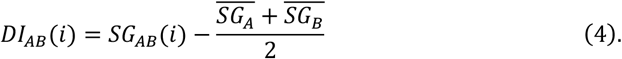

If the plant-growth promoting effects under the presence of two fungal isolates is close to what expected as the intermediate results of the single inoculation assays of the two isolates, the index for deviation from intermediate effects [*DI*_*AB*_(*i*)] or its mean over replicate samples 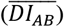 is likely to have a value around zero.

### Nonlinearity of fungus–fungus combinations

For each pair of fungal isolates (A and B), an analysis of variance (ANOVA) model of standardized growth index was constructed by including the presence/absence of isolate A, the presence/absence of isolate B, and the interaction term of the two (i.e., isolate A × isolate B) as explanatory variables. Then, across the 78 fungal pairs examined, *F* values of the isolate A × isolate B term were compared as indicators of how combinations of the two fungal isolates had “nonlinear” effects on plant performance traits. We then examined how the nonlinearity measures of fungal pairs are associated with the abovementioned index values representing deviations of observed dual-inoculation results from those expected as intermediate results of single inoculations 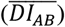.

All the calculations of the above indexes and statistical analyses were performed using the R ver. 3.6.0.

## RESULTS

### Germination rates

The gemination rates of *Brassica* plants varied within single inoculation treatments and within dual inoculation treatments (Supplementary Figure S2). Meanwhile, the rates were generally higher in dual inoculation treatments than in single inoculation treatments (Welch’s test; *t* = -3.97, df = 13.6, *P* = 0015).

### Plants’ growth responses

For all the three plant performance variables (shoot dry weight, the number of mature leaves, and SPAD), the single inoculation effects on *Brassica* plants differed significantly among the 13 fungal isolates examined (Table 2). For example, the mean standardized growth index for *V. simplex* Y34 and *Alternaria* sp. KYOCER00001239 were, on average, ca. seven-fold larger than the standard deviation of control sample’s growth (i.e., 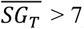) in terms of shoot dry weight, indicating high growth-promoting effects of those fungi on *Brassica* plants (Fig. 2A). In addition, *C. chaetospira* M4006, *Trichoderma* sp. KYOCER00000218, *Curvularia* sp. KYOCER00000077, *Phoma* sp. KYOCER00000052, and *Stemphylium* sp. KYOCER00000804 showed high plant growth promoting effects (Fig. 2A). In contrast, *C. tofieldiae* MAFF 712334, *Mucor* sp. KYOCER00000113, *Setophaeria* sp. KYOCER00000031, *Fusarium* sp. KYOCER00000983 and *Tolypocladium* sp. KYOCER00000289 displayed weak or almost neutral effects on plant growth and *Aspergillus* sp. KYOCER00000917 had negative impacts on the *Brassica* plants (Fig. 2A). When the number of mature leaves was used as a metric of plant performance, *Alternaria* sp. KYOCER00001239 and *Aspergillus* sp. KYOCER00000917 turned out to have strongly positive and negative effects, respectively (Fig. 2B). Meanwhile, the effects of other fungal isolates were moderately positive or neutral (Fig. 2B).

**TABLE 2.** ANOVA results of single- and dual-inoculation experiments.

| ANOVA model | df | *F* | *P* |
| --- | --- | --- | --- |
| Single inoculation (across 13 fungal isolates) |  |  |  |
| Shoot dry weight | 12 | 41.6 | < 0.0001 |
| Number of mature leaves | 12 | 127.7 | < 0.0001 |
| SPAD readings | 12 | 12.0 | < 0.0001 |
| Dual inoculation (across 78 fungal pairs) |  |  |  |
| Shoot dry weight | 77 | 25.7 | < 0.0001 |
| Number of mature leaves | 77 | 23.1 | < 0.0001 |
| SPAD readings | 77 | 5.7 | < 0.0001 |

For each of the three plant performance variables, an ANOVA model was constructed to examine the variation across single- or dual-inoculation treatments.

**FIGURE 1.**
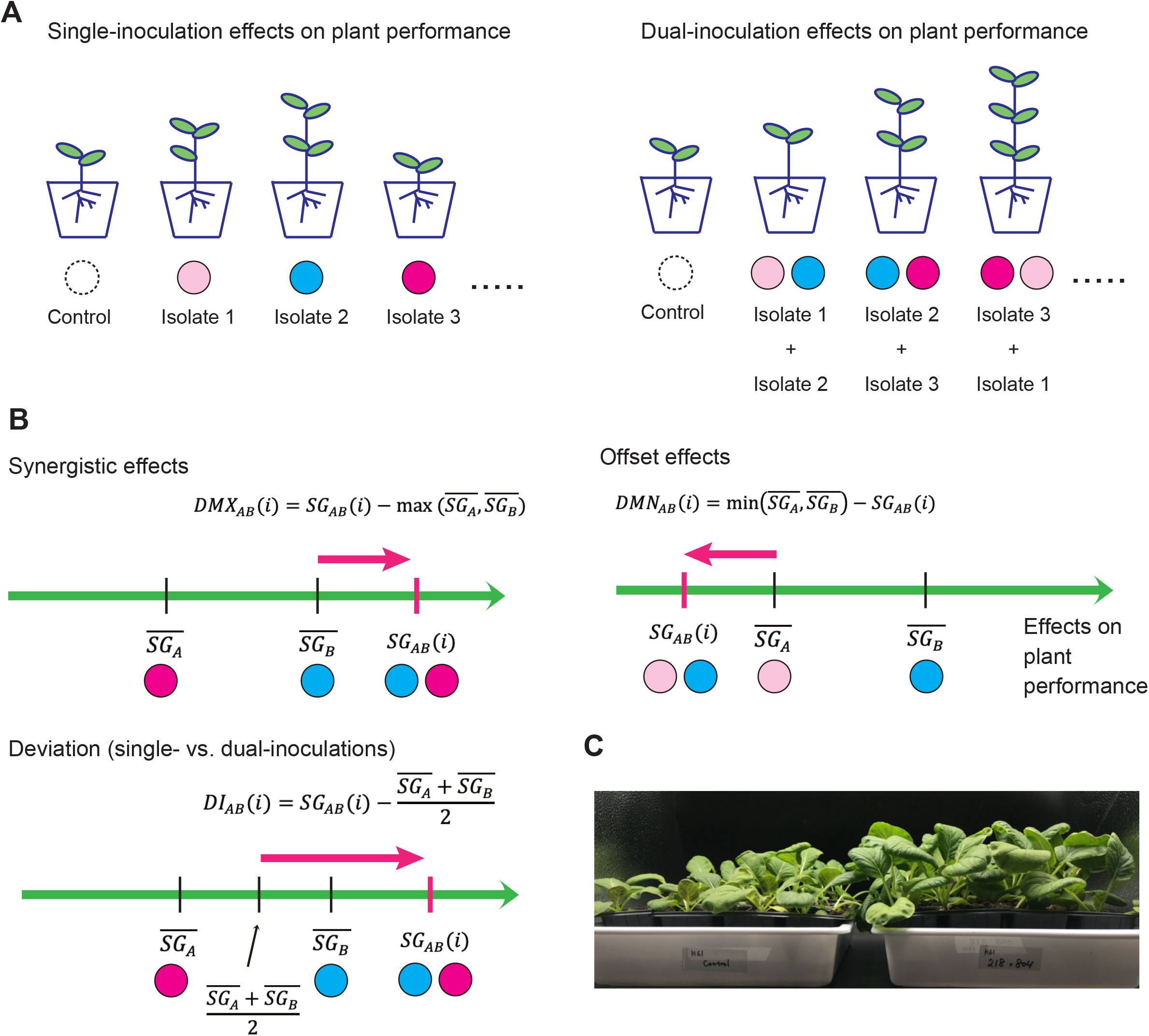
Evaluation of effects on plant performance. **(A)** Schema of single- and dual-inoculation assays. **(B)** Indexes for comparing single vs. dual inoculation effects. Along the axis of standardized growth index defined by the equation (1), index values representing synergistic/offset effects on plants are calculated for each replicate plant sample for each pair of microbial (fungal) isolates [*DMX*_*AB*_(*i*) and *DMN*_*AB*_(*i*)]. Likewise, index values representing deviation of dual-inoculation effects from single-inoculation effects are obtained [*DI*_*AB*_(*i*)]. **(C)** Example of inoculation experiments. *Brassica* plants inoculated with two fungal isolates (tri_0218 × ste_0804; right) and those without fungal inoculations (control; left).

**FIGURE 2.**
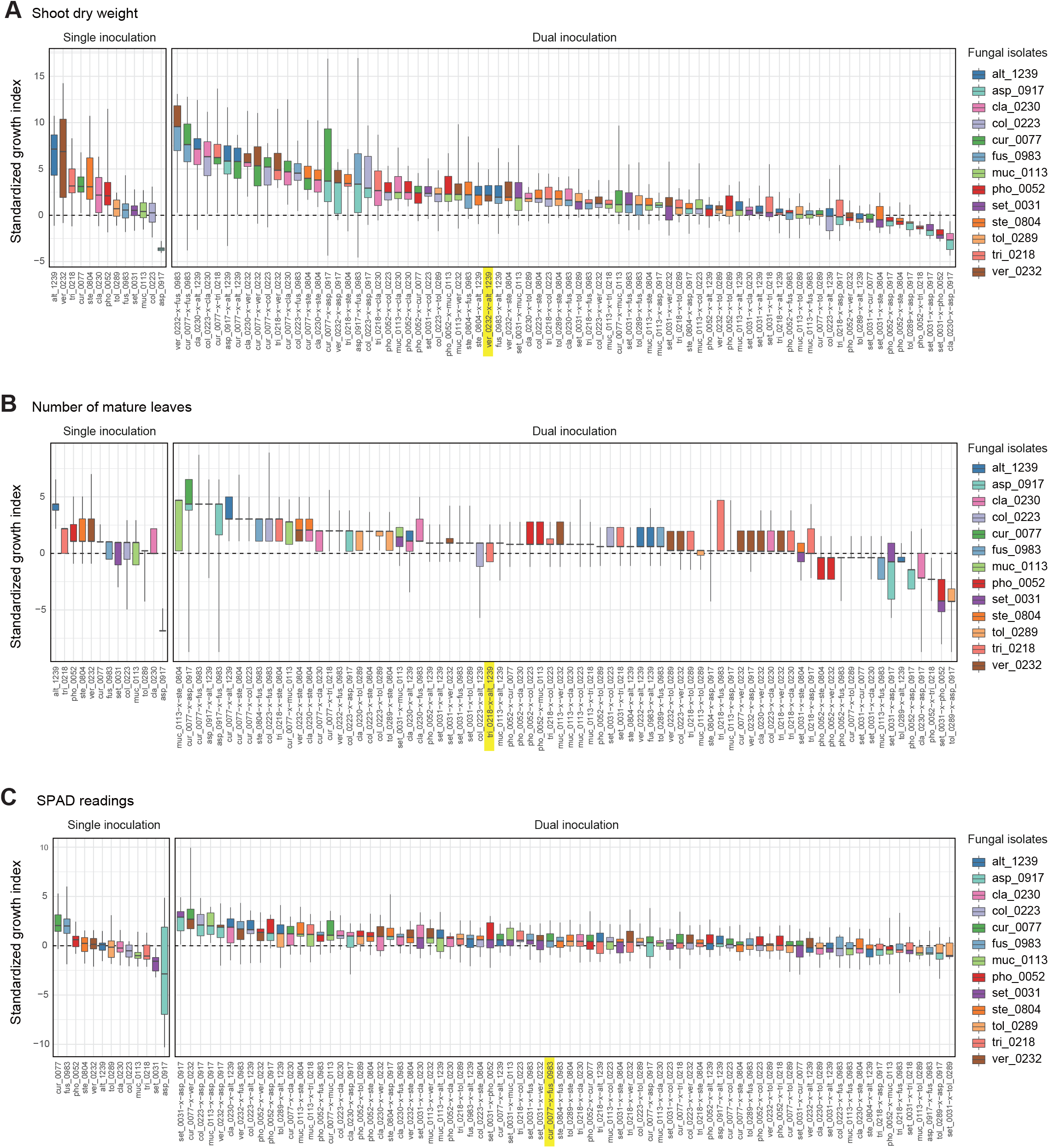
Single- and dual-inoculation effects on *Brassica* plants. **(A)** Standardized growth index in terms of shoot dry weight. For respective single- and dual-inoculation experiments, 25 % quantiles, medians, and 75 % quantiles are displayed as boxes and the ranges from the maximum to minimum values are shown as bars. See Table 1 for the abbreviation of fungal isolates. The combination of the fungal species with the largest positive effects on *Brassica* plants in single inoculation experiments is highlighted. **(B)** Standardized growth index in terms of the number of mature leaves. **(C)** Standardized growth index in terms of SPAD readings.

In the dual inoculation experiments, the pair of the fungal isolates that exhibited the greatest effects in single inoculation treatments (i.e., *V. simplex* Y34 and *Alternaria* sp. KYOCER00001239) had relatively weak positive effects on *Brassica* growth in terms of shoot dry weight (Fig. 2A). Instead, the highest plant-growth promoting effects were observed for the combination of *V. simplex* Y34 and *Fusarium* sp. KYOCER00000983, which had neutral effects on plants in the single inoculation (Fig. 2A). Highly positive effects on plants (e.g., 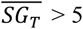) were observed, as well, in *Curvularia–Fusarium, Cladophialophora*– *Alternaria, Colletotrichum*–*Cladophialophora, Aspergillus*–*Alternaria*, and *Cladophialophora*–*Veronaeopsis* pairs and several other pairs including *Curvularia* sp. KYOCER00000077: for these pairs, at least one partner had neutral to weakly positive performance in single inoculation treatments (Fig. 2A).

In contrast to those combinations with relatively high plant-growth promoting effects (in the metrics of shoot dry weight and the number of mature leaves), *Aspergillus* sp. KYOCER00000917, which restricted plant growth in the single inoculation condition (Fig. 2A, B), had negative impacts on plants in some of the 12 combinations with other fungal isolates (Fig. 3A, B). However, their negative effects diminished in dual inoculations with some fungi such as *Alternaria* sp. KYOCER00001239 and *Curvularia* sp. KYOCER00000077 (Fig. 3A, B). Results also showed that *Phoma* sp. KYOCER00000052, whose impacts on plants were positive in the single inoculation setting, inhibited plant growth in the presence of other fungi (Fig. 3A, B).

**FIGURE 3.**
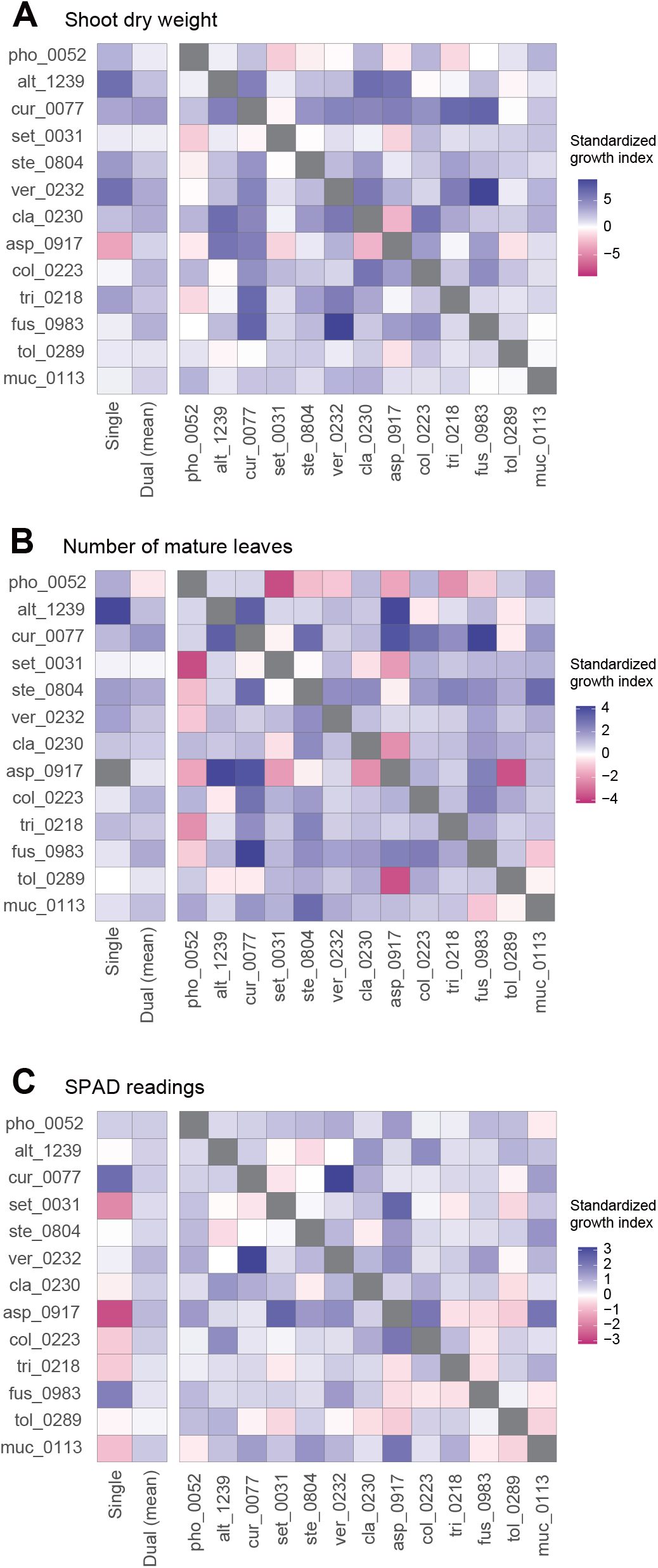
Pairwise representation of dual inoculation results. **(A)** Standardized growth index in terms of shoot dry weight for each pair of fungal isolates. Single-inoculation effects and mean effects across the dual inoculation assays are shown for each fungal isolate in the left. **(B)** Standardized growth index in terms of the number of mature leaves. **(C)** Standardized growth index in terms of SPAD readings.

When SAPD readings were used as metrics of plant performance, the *Curvularia* sp. KYOCER00000077 and *Fusarium* sp. KYOCER00000983 had relatively high positive effects on *Brassica* plants 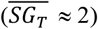, while *Setophaeria* sp. KYOCER00000031 and *Aspergillus* sp. KYOCER00000917 had negative impacts (Fig. 2C). Note that SPAD readings were weakly correlated with shoot dry weight and the number of mature leaves (Supplementary Fig. S3). In the dual inoculation experiments, some fungal pairs including *Aspergillus* sp. KYOCER00000917 had relatively high positive effects on *Brassica* plants (Fig. 3C) despite negative impacts of the *Aspergillus* isolate in a single-inoculation condition (Fig. 2C). The pair of *Curvularia* and *Veronaeopsis* moderately increased SPAD readings as well (Fig. 3C). Meanwhile, SPAD readings did not differ greatly from the control for most fungal pairs (Fig. 2C).

For all the three plant performance variables examined, standardized growth index values of single inoculation experiments were uncorrelated with those averaged across dual inoculations for respective fungi (shoot dry weight, *r* = -0.09, *P* = 0.78; number of mature leaves *r* = 0.11, *P* = 0.71; SPAD, *r* = -0.41, *P* = 0.17; Fig. 3). In other words, fungi with more positive effects on plants in single-inoculation experiments did not increased plant performance more efficiently. The experimental results also indicated that some combinations of fungi exhibited higher impacts on *Brassica* performance than that observed in all the single-inoculation settings (Fig. 2A-C).

### Synergistic and offset effects

Among the 78 combinations of fungal isolates, strong synergistic effects 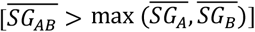 were observed in some pairs of fungi in terms of shoot dry weight (Fig. 4A). The fungal combinations with the largest synergistic effects 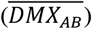 consisted of *Curvularia* sp. KYOCER00000077 and *Fusarium* sp. KYOCER00000983, each of which had weakly or moderately positive impacts on plant growth in single inoculations. Large synergistic effects were detected in other pairs of fungi including fungi with moderate or weakly positive effects on plants (e.g., *Colletotrichum–Cladophialophora, Colletotrichum– Fusarium*, and *Veronaeopsis–Fusarium* pairs; Fig. 4A). Similarly, for the number of mature leaves, fungal pairs with large synergistic effects involved fungi with weakly positive or even negative effects in single inoculations (Fig. 4B). In terms of SPAD readings, pairs of fungi with negative impacts on plants in single-inoculation conditions had large synergistic effects (Fig. 4C).

**FIGURE 4.**
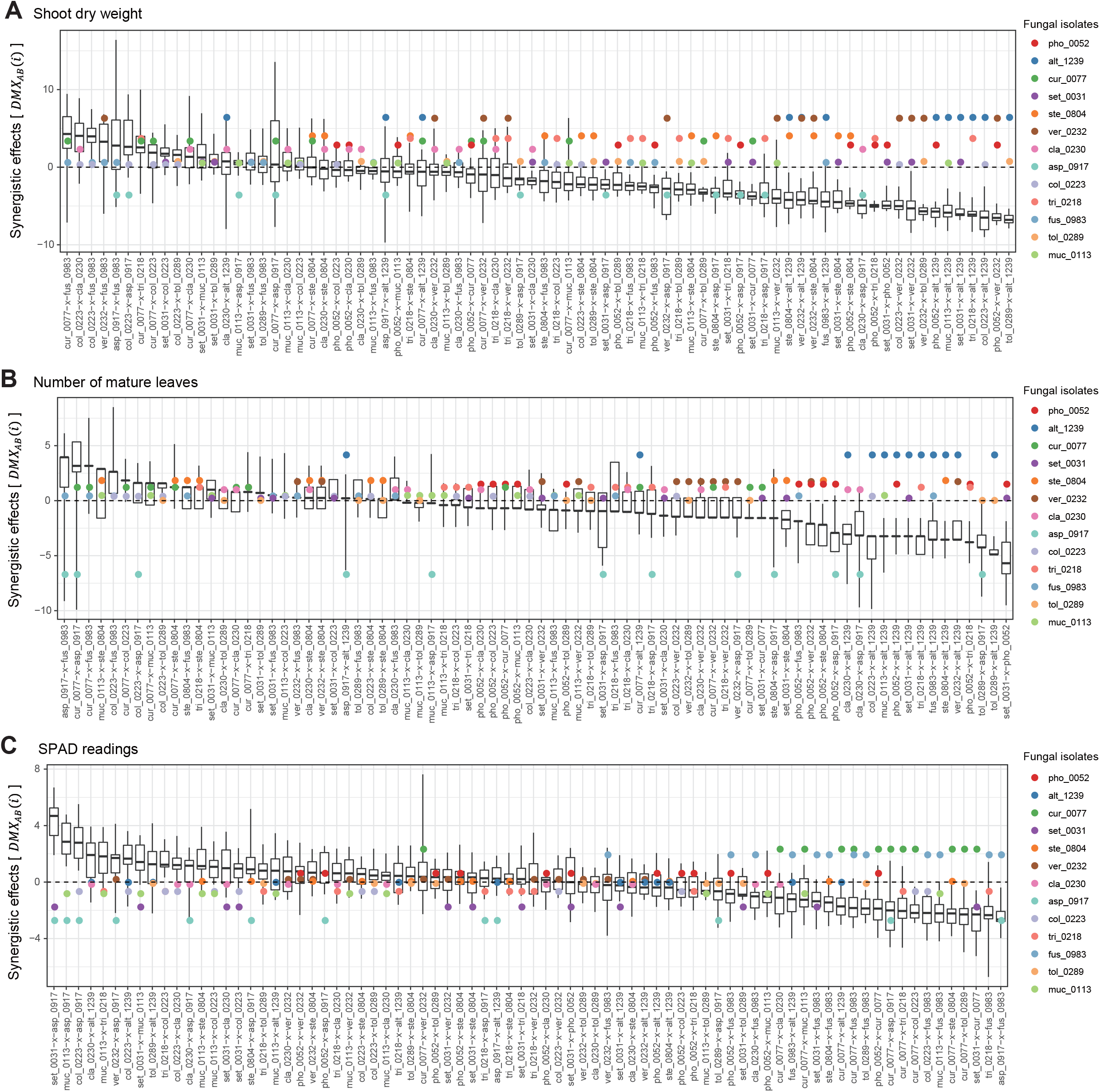
Synergistic effects observed in dual-inoculation experiments. **(A)** Synergistic effect index in terms of shoot dry weight. The index representing deviation of dual-inoculation effects from the maximum effects in single inoculations are shown for each pair of fungal isolates. Circles represent single-inoculation effects of respective fungal isolates. **(B)** Synergistic effect index in terms of the number of mature leaves. **(C)** Synergistic effect index in terms of SPAD readings.

In contrast to synergistic effects, offset effects 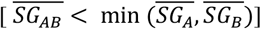 were evident especially in the fungal pairs including fungi that had highly positive impacts on plant performance traits in single-inoculation conditions (Fig. 5). In particular, the pairs of fungi with the largest positive effects (i.e., the *Veronaeopsis–Alternaria* pair) showed large offset effects (Fig. 5).

**FIGURE 5.**
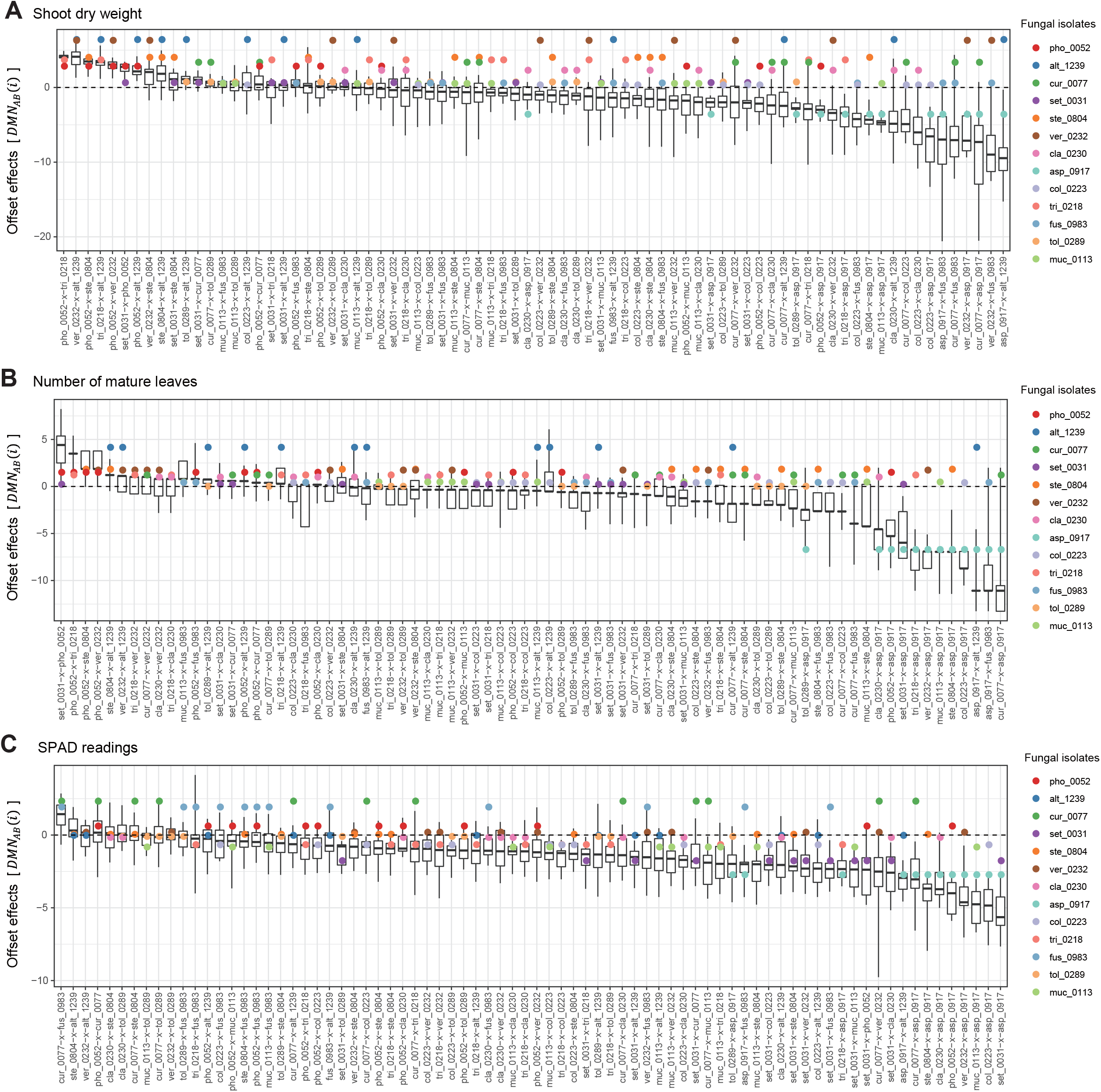
Offset effects observed in dual-inoculation experiments. **(A)** Offset effect index in terms of shoot dry weight. The index representing deviation of dual-inoculation effects from the minimum effects in single inoculations are shown for each pair of fungal isolates. Circles represent single-inoculation effects of respective fungal isolates. **(B)** Offset effect index in terms of the number of mature leaves. **(C)** Offset effect index in terms of SPAD readings.

Across the 78 combinations of fungi, synergistic effects (i.e., 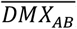) decreased with increasing mean values of single inoculation effects of the target fungi (i.e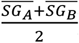) (Fig. 6A-C). In other words, pairs of fungi that showed greater plant-performance increasing effects tended to have weaker synergistic effects. As expected by the trend in synergistic effects, offset effects were increased with increasing mean values of single inoculation effects of the target fungi (Fig. 6D-F).

**FIGURE 6.**
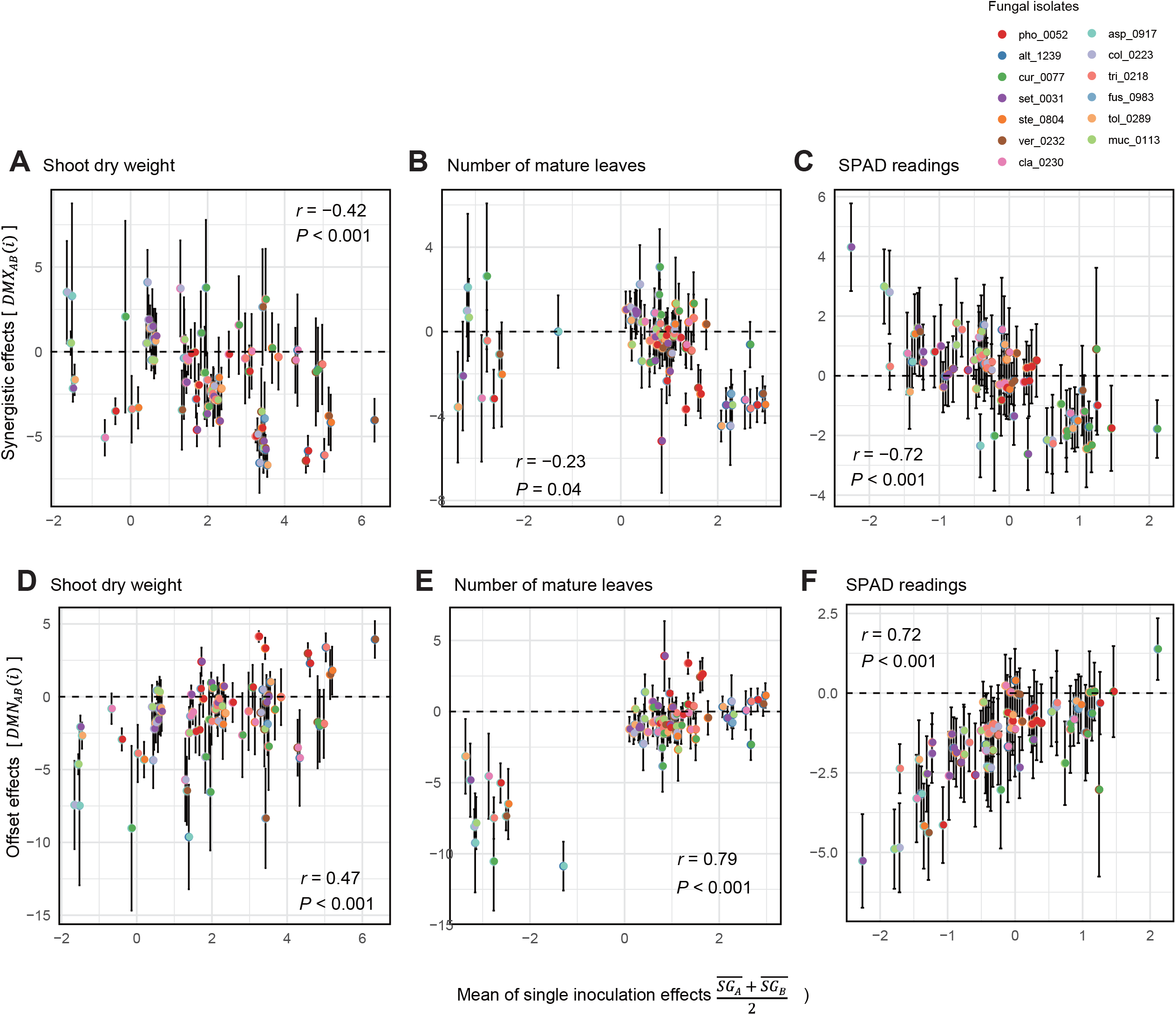
Relationship between single-inoculation effects and synergistic/offset effects. **(A)** Trends in synergistic effects in terms of shoot dry weight. For each pair of fungi, mean values of single inoculation effects of the target fungi (i.e., 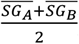) and index values of synergistic effects [i.e., *DMX*_*AB*_(*i*)] are shown at the horizontal and vertical axes, respectively. Error bars represent standard deviations of synergistic effects. **(B)** Trends in synergistic effects in terms of the number of mature leaves. **(C)** Trends in synergistic effects in terms of SPAD readings. **(D)** Trends in synergistic effects in terms of shoot dry weight. For each pair of fungi, mean values of single inoculation effects of the target fungi (i.e., 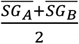) and index values of offset effects [i.e., *DMN*_*AB*_(*i*)] are shown at the horizontal and vertical axes, respectively. **(E)** Trends in synergistic effects in terms of the number of mature leaves. **(F)** Trends in synergistic effects in terms of SPAD readings.

### Nonlinearity of fungus–fungus combinations

Deviations of observed dual-inoculation results from those expected as intermediate results of single inoculations 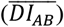 varied among fungal pairs (Fig. 6). Higher absolute 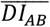 values were indicative of nonlinearity in effects on plants for the particular fungus–fungus combinations as evaluated by a series of ANOVA models (Fig. 7; Supplementary Data S3).

**FIGURE 7.**
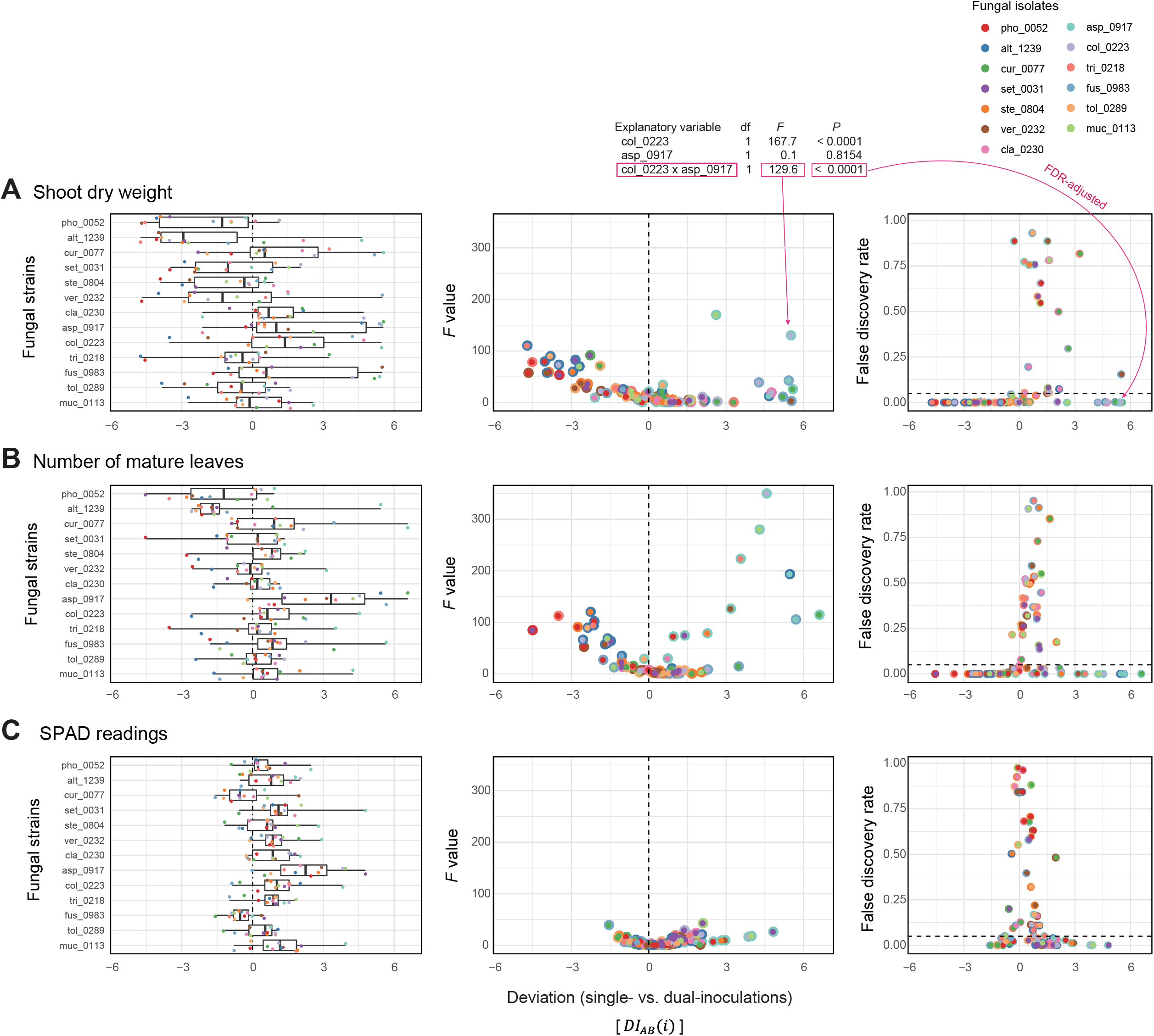
Deviations of observed dual-inoculation results from those expected as intermediate results of single inoculations. **(A)** Deviation index for shoot dry weight. The index values representing deviations of dual-inoculation effects from intermediate effects in single inoculations 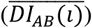 are shown for each fungal isolate included in the target fungal pairs (left). For each fungal pair, *F* values of the isolate A × isolate B term in the ANOVA model (middle) and false discovery rate (FDR) values of the interaction term (right) are shown across the axis of the deviation index: FDR are calculated across the 78 fungal combinations examined. **(B)** Deviation index for the number of mature leaves. **(C)** Deviation index for SPAD readings.

## DISCUSSION

By using taxonomically diverse endophytic and soil fungi, we here evaluated plant-growth promoting effects of pairs of fungal isolates in light of those observed in single-isolate inoculation experiments. The 13 fungal isolates differed greatly in their independent effects on *Brassica* plants (Figs. 2-3), providing an ideal opportunity for examining how the ranking of plant-growth promoting effects in single-inoculation contexts were related to that in multi-species (dual-inoculation) contexts (Figs. 4-6). Such information of synergistic and offset effects in the presence of multiple microbial species is indispensable for understanding to what extent we can predict functions of microbial communities (microbiomes) from the datasets of single-species/isolate screening.

A series of single- and dual-inoculation experiments indicated that greater performance of plants are potentially obtained in multi-species than in single-species contexts (Fig. 2). This result, itself, is consistent with previous reports of enhanced plant growth by specific pairs of bacteria/fungi (Han and Lee, 2006; Wang et al., 2011; Ważny et al., 2018; He et al., 2020). Meanwhile, our experiments on 78 combinations of fungi suggested that pairs of microbes, each of which had greatly positive impacts on plant growth in single inoculations, could show minor effects on plants in multi-species conditions. For example, the strategy of combining the two “highest rankers” in the single inoculation experiments (i.e., *Veronaeopsis simplex* and *Alternaria* sp. KYOCER00001239) did not result in high plant-growth promoting effects (Fig. 3): rather, offset effects were observed in the highest ranker pairs (Figs. 4-6). Thus, biological functions at the community (microbiome) level may be rarely maximized by the “bottom-up” exploration of sets of microbes based solely on single-inoculation experiments.

Our experiments also suggested that pairs of microbes with subordinate performance in single inoculation assays could show largest growth-promoting effects on plants (Fig. 2). This result suggests that single-species/isolate screening does not always provide sufficient information for predicting microbial performance at the multi-species level (Toju et al., 2018a). Interestingly, the fungal pairs with highest synergistic effects in our experiment involved fungi in the genera *Fusarium* and *Curvularia* (Fig. 4A), which were often described as plant pathogenic taxa (Michielse and Rep, 2009; Ma et al., 2013; Manamgoda et al., 2015). Basically, physiological effects on plants vary remarkably among species/isolates within taxa as evidenced by the presence of *Fusarium* and *Curvularia* species enhancing plant health and growth (Olivain et al., 2006; Nahalkova et al., 2008; Priyadharsini and Muthukumar, 2017).

In fact, the *Fusarium* and *Curvularia* isolates examined in our study had positive effects on *Brassica* plants even in the single-inoculation assays (Fig. 2). Moreover, the results of the dual inoculation experiments suggested that some fungi in these predominantly plant-pathogenic genera can have even greater positive effects on plants in combination with specific other fungi (Figs. 2-3). Our results on synergistic effects in multi-species contexts further illuminate the potential use of diverse endosphere/rhizosphere microbes whose biological functions have been underestimated in conventional screening of single inoculations.

The fact that microbial functions critically depend on combinations of microbial species/isolates highlight the importance of “bird’s-eye” views of designing microbiomes. Given that microbial functions at the community (multi-species) levels are not the simple sums/averages of functions in single-species contexts (Figs. 2 & 7), research strategies taking into account not only each microbe’s roles but also the nature of microbe–microbe interactions will provide platforms for optimization of microbiome functions (Agler et al., 2016; Toju et al., 2016; Banerjee et al., 2018). In this respect, interdisciplinary approaches integrating the observational, genomic, and metagenomic information of microbial functions (Bulgarelli et al., 2015; Levy et al., 2018; Ichihashi et al., 2020) with community ecological analyses of species interaction networks (Agler et al., 2016; van der Heijden and Hartmann, 2016; Toju et al., 2017) will help us explore highly functional and stable microbial sets among numerous candidate combinations of species (Paredes et al., 2018; Saad et al., 2020; Toju et al., 2020). In other words, information of microbial functions in single-species contexts is utilized by being combined with insights into dynamics and processes within microbiomes.

While the experiments conducted in this study provided a unique opportunity for systematically evaluating synergistic/offset effects of microbes on plants, the obtained datasets should be interpreted with caution given the following limitations. First, physiological mechanisms by which the examined fungi affected plant growth were unexplored in the current study. Although detailed physiological and/or molecular biological investigations have been done for some of the fungal species used in this study [e.g., *C. tofieldiae* (Hiruma et al., 2016), *Veronaeopsis simplex* (Guo et al., 2018), and *C. chaetospira* (Harsonowati et al., 2020)], metabolites and genes involved in the plant–fungus interactions are unknown for the remaining species. For more mechanistic understanding of interactions involving plants and multiple microbial species, we need to perform transcriptomic analyses targeting plants’ responses to each microbe as well as those comparing plants’ gene expression patterns between single- and multiple-symbiont conditions. Comparative transcriptomic analyses across experiments with different environmental conditions (e.g., soil nutrient concentrations) will provide essential insights into microbial functions as well. Second, the inoculation test based on single plant species precluded us from understanding how general synergistic/offset effects existed in plant–fungal biome interactions. Although some of the fungal taxa used in this study have been reported to interact with multiple families of plants (Hermosa et al., 2012; Toju et al., 2018b), impacts of endophytic/soil fungi on plants can vary depending on plant taxa and environmental conditions (Kiers et al., 2011; Pineda et al., 2013; Rudgers et al., 2020). Therefore, to gain more robust insights into synergistic/offset effects in interactions of plants and multiple microbial species/isolates, the reproducibility of the patterns observed in this study should be examined in inoculation experiments targeting diverse other plant species. Third, it is important to acknowledge that the complexity of the microbial sets examined in this study is minimal (i.e., two fungal species): different types of phenomena may be observed in combinations of three or more bacterial/fungal species (Durán et al., 2018; Paredes et al., 2018; Carlström et al., 2019; Wei et al., 2019)(Durán et al., 2018; Paredes et al., 2018; Carlström et al., 2019; Wei et al., 2019). Moreover, it remains to be examined how we can increase microbial functions (e.g., host plant growth rates) by increasing the number of microbial species/isolates. The presence of microbial pairs outperforming single-microbe systems (Fig. 2) leads to the working hypothesis that compatible sets of three or more microbial species yield greater functions than simpler communities by playing complementary roles. Meanwhile, it is expected that benefits of microbiomes do not increase linearly with increasing number of microbial species (i.e., saturating curves of benefits against increasing number of microbes) (van der Heijden et al., 1998), at least in terms of specific functions such as provisioning of soil phosphorus or blocking of soil pathogens.

We here showed that screening based on inoculations of single microbial species/isolates can result in the underestimation of the microbes that potentially have large plant-growth promoting effects in combinations with specific other microbes. Given that plants are inevitably associated with hundreds or more of microbial species in agricultural and natural ecosystems (Lundberg et al., 2012; Schlaeppi and Bulgarelli, 2015; van der Heijden and Hartmann, 2016), such nonlinearity found in microbe–microbe associations deserve future intensive research. Interdisciplinary studies on relationships between microbiome compositions and their ecosystem-level functions are awaited towards the maximization of microbial functions for sustainable agriculture and ecosystem restoration.

## AUTHOR CONTRIBUTIONS

YH and HT designed the work. YH carried out the experiments. YH, HF, and HT analyzed the data. YH and HT wrote the manuscript based on discussion with HF, KH, and KN.

## ACKNOWLEDGEMENTS

This work was financially supported in part by Precursory Research for Embryonic Science and Technology (PRESTO) of Japan Science and Technology Agency (JST; JPMJPR16Q6 to HT), Human Frontier Science Program (HFSP; RGP0029/2019 to HT), and New Energy and Industrial Technology Development Organization (NEDO; JPNP18016 to HT).

## SUPPLEEMENTARY MATERIAL

The Supplementary Material for this article can be found online at XXXXX.

## REFERENCES

Agler, M. T., Ruhe, J., Kroll, S., Morhenn, C., Kim, S. T., Weigel, D., et al. (2016). Microbial hub taxa link host and abiotic factors to plant microbiome variation. PLoS Biology 14, 1–31. doi:10.1371/journal.pbio.1002352.

Ahmad, F., Ahmad, I., and Khan, M. S. (2008). Screening of free-living rhizospheric bacteria for their multiple plant growth promoting activities. Microbiological Research 163, 173–181. doi:10.1016/j.micres.2006.04.001.

Banerjee, S., Schlaeppi, K., and van der Heijden, M. G. A. (2018). Keystone taxa as drivers of microbiome structure and functioning. Nature Reviews Microbiology 16, 567–576. doi:10.1038/s41579-018-0024-1.

Bhattacharyya, P. N., and Jha, D. K. (2012). Plant growth-promoting rhizobacteria (PGPR): Emergence in agriculture. World Journal of Microbiology and Biotechnology 28, 1327–1350. doi:10.1007/s11274-011-0979-9.

Bulgarelli, D., Garrido-Oter, R., Münch, P. C., Weiman, A., Dröge, J., Pan, Y., et al. (2015). Structure and function of the bacterial root microbiota in wild and domesticated barley. Cell Host and Microbe 17, 392–403. doi:10.1016/j.chom.2015.01.011.

Bulgarelli, D., Schlaeppi, K., Spaepen, S., van Themaat, E. V. L., and Schulze-Lefert, P. (2013). Structure and functions of the bacterial microbiota of plants. Annual Review of Plant Biology 64, 807–838. doi:10.1146/annurev-arplant-050312-120106.

Busby, P. E., Soman, C., Wagner, M. R., Friesen, M. L., Kremer, J., Bennett, A., et al. (2017). Research priorities for harnessing plant microbiomes in sustainable agriculture. PLoS Biology 15, 1–14. doi:10.1371/journal.pbio.2001793.

Carlström, C. I., Field, C. M., Bortfeld-Miller, M., Müller, B., Sunagawa, S., and Vorholt, J. A. (2019). Synthetic microbiota reveal priority effects and keystone strains in the Arabidopsis phyllosphere. Nature Ecology and Evolution 3, 1445–1454. doi:10.1038/s41559-019-0994-z.

Castrillo, G., Teixeira, P. J. P. L., Paredes, S. H., Law, T. F., de Lorenzo, L., Feltcher, M. E., et al. (2017). Root microbiota drive direct integration of phosphate stress and immunity. Nature 543, 513–518. doi:10.1038/nature21417.

Chang, S. X., and Robison, D. J. (2003). Nondestructive and rapid estimation of hardwood foliar nitrogen status using the SPAD-502 chlorophyll meter. Forest Ecology and Management 181, 331–338. doi:10.1016/S0378-1127(03)00004-5.

Dai, C. C., Yu, B. Y., and Li, X. (2008). Screening of endophytic fungi that promote the growth of Euphorbia pekinensis. African Journal of Biotechnology 7, 3505–3510. doi:10.4314/ajb.v7i19.59361.

Durán, P., Thiergart, T., Garrido-Oter, R., Agler, M., Kemen, E., Schulze-Lefert, P., et al. (2018). Microbial interkingdom interactions in roots promote Arabidopsis survival. Cell 175, 973–983. doi:10.1016/j.cell.2018.10.020.

Esfahani, M., Abbasi, H. R. A., Rabiei, B., and Kavousi, M. (2008). Improvement of nitrogen management in rice paddy fields using chlorophyll meter (SPAD). Paddy and Water Environment 6, 181–188. doi:10.1007/s10333-007-0094-6.

Finkel, O. M., Salas-González, I., Castrillo, G., Conway, J. M., Law, T. F., Teixeira, P. J. P. L., et al. (2020). A single bacterial genus maintains root growth in a complex microbiome. Nature 587, 103–108. doi:10.1038/s41586-020-2778-7.

Gu, S., Wei, Z., Shao, Z., Friman, V. P., Cao, K., Yang, T., et al. (2020). Competition for iron drives phytopathogen control by natural rhizosphere microbiomes. Nature Microbiology 5, 1002–1010. doi:10.1038/s41564-020-0719-8.

Guo, Y., Matsuoka, Y., Nishizawa, T., Ohta, H., and Narisawa, K. (2018). Effects of rhizobium species living with the dark septate endophytic fungus veronaeopsis simplex on organic substrate utilization by the host. Microbes and Environments 33, 102–106. doi:10.1264/jsme2.ME17144.

Han, H. S., and Lee, K. D. (2006). Effect of co-inoculation with phosphate and potassium solubilizing bacteria on mineral uptake and growth of pepper and cucumber. Plant, Soil and Environments 52, 130–136.

Harbort, C. J., Hashimoto, M., Inoue, H., Niu, Y., Guan, R., Rombolà, A. D., et al. (2020). Root-secreted coumarins and the microbiota interact to improve iron nutrition in Arabidopsis. Cell Host and Microbe 28, 825–837. doi:10.1016/j.chom.2020.09.006.

Harsonowati, W., Marian, M., Surono, M., and Narisawa, K. (2020). The effectiveness of a dark septate endophytic fungus, Cladophialophora chaetospira SK51, to mitigate strawberry Fusarium wilt disease and with growth promotion activities. Frontiers in Microbiology 11, 585–585. doi:10.3389/fmicb.2020.00585.

He, C., Wang, W., and Hou, J. (2020). Plant performance of enhancing licorice with dual inoculating dark septate endophytes and Trichoderma viride mediated via effects on root development. BMC Plant Biology 20, 1–14. doi:10.1186/s12870-020-02535-9.

Helfrich, E. J. N., Vogel, C. M., Ueoka, R., Schäfer, M., Ryffel, F., Müller, D. B., et al. (2018). Bipartite interactions, antibiotic production and biosynthetic potential of the Arabidopsis leaf microbiome. Nature Microbiology 3, 909–919. doi:10.1038/s41564-018-0200-0.

Hermosa, R., Viterbo, A., Chet, I., and Monte, E. (2012). Plant-beneficial effects of Trichoderma and of its genes. Microbiology 158, 17–25. doi:10.1099/mic.0.052274-0.

Hiruma, K., Gerlach, N., Sacristán, S., Nakano, R. T., Hacquard, S., Kracher, B., et al. (2016). Root Endophyte Colletotrichum tofieldiae Confers Plant Fitness Benefits that Are Phosphate Status Dependent. Cell 165, 464–474. doi:10.1016/j.cell.2016.02.028.

Hiruma, K., Kobae, Y., and Toju, H. (2018). Beneficial associations between Brassicaceae plants and fungal endophytes under nutrient-limiting conditions : evolutionary origins and host –symbiont molecular mechanisms. Current Opinion in Plant Biology 44, 145–154. doi:10.1016/j.pbi.2018.04.009.

Ichihashi, Y., Ichihashi, Y., Date, Y., Date, Y., Shino, A., Shimizu, T., et al. (2020). Multiomics analysis on an agroecosystem reveals the significant role of organic nitrogen to increase agricultural crop yield. Proceedings of the National Academy of Sciences of the United States of America 117, 14552–14560. doi:10.1073/pnas.1917259117.

Jansa, J., Forczek, S. T., Rozmoš, M., Püschel, D., Bukovská, P., and Hršelová, H. (2019). Arbuscular mycorrhiza and soil organic nitrogen: network of players and interactions. Chemical and Biological Technologies in Agriculture 6, 1–10. doi:10.1186/s40538-019-0147-2.

Kennedy, P. G., Peay, K. G., and Bruns, T. D. (2009). Root tip competition among ectomycorrhizal fungi: are priority effects a rule or an exception? Ecology 90, 2098–107. Available at: http://www.ncbi.nlm.nih.gov/pubmed/19739372.

Khastini, R. O., Ohta, H., and Narisawa, K. (2012). The Role of a Dark Septate Endophytic Fungus, Veronaeopsis simplex Y34, in Fusarium Disease Suppression in Chinese Cabbage. 50, 618–624. doi:10.1007/s12275-012-2105-6.

Kiers, E. T., Duhamel, M., Beesetty, Y., Mensah, J. A., Franken, O., Verbruggen, E., et al. (2011). Reciprocal rewards stabilize cooperation in the mycorrhizal symbiosis. Science 333, 880–882. doi:10.1126/science.1208473.

Levy, A., Salas Gonzalez, I., Mittelviefhaus, M., Clingenpeel, S., Herrera Paredes, S., Miao, J., et al. (2018). Genomic features of bacterial adaptation to plants. Nature Genetics 50, 138–150. doi:10.1038/s41588-017-0012-9.

Lugtenberg, B., and Kamilova, F. (2009). Plant-growth-promoting rhizobacteria. Annual Review of Microbiology 63, 541–556. doi:10.1146/annurev.micro.62.081307.162918.

Lundberg, D. S., Lebeis, S. L., Paredes, S. H., Yourstone, S., Gehring, J., Malfatti, S., et al. (2012). Defining the core Arabidopsis thaliana root microbiome. Nature 488, 86–90. doi:10.1038/nature11237.

Ma, L. J., Geiser, D. M., Proctor, R. H., Rooney, A. P., O’Donnell, K., Trail, F., et al. (2013). Fusarium pathogenomics. Annual Review of Microbiology 67, 399–416. doi:10.1146/annurev-micro-092412-155650.

Manamgoda, D. S., Rossman, A. Y., Castlebury, L. A., Chukeatirote, E., and Hyde, K. D. (2015). A taxonomic and phylogenetic re-appraisal of the genus Curvularia (Pleosporaceae): Human and plant pathogens. Phytotaxa 212, 175–198. doi:10.11646/phytotaxa.212.3.1.

Michielse, C. B., and Rep, M. (2009). Pathogen profile update: Fusarium oxysporum. Molecular Plant Pathology 10, 311–324. doi:10.1111/j.1364-3703.2009.00538.x.

Nahalkova, J., Fatehi, J., Olivain, C., and Alabouvette, C. (2008). Tomato root colonization by fluorescent-tagged pathogenic and protective strains of Fusarium oxysporum in hydroponic culture differs from root colonization in soil. FEMS Microbiology Letters 286, 152–157. doi:10.1111/j.1574-6968.2008.01241.x.

Nara, K. (2006). Ectomycorrhizal networks and seedling establishment during early primary succession. New Phytologist 169, 169–178. doi:10.1111/j.1469-8137.2005.01545.x.

Narisawa, K., Usuki, F., and Hashiba, T. (2004). Control of Verticillium yellows in Chinese cabbage by the dark septate endophytic fungus LtVB3. Phytopathology 94, 412–418. doi:10.1094/PHYTO.2004.94.5.412.

Nelson, J. M., Hauser, D. A., Hinson, R., and Shaw, A. J. (2018). A novel experimental system using the liverwort Marchantia polymorpha and its fungal endophytes reveals diverse and context-dependent effects. New Phytologist 218, 1217–1232. doi:10.1111/nph.15012.

Netto, A. T., Campostrini, E., de Oliveira, J. G., and Bressan-Smith, R. E. (2005). Photosynthetic pigments, nitrogen, chlorophyll a fluorescence and SPAD-502 readings in coffee leaves. Scientia Horticulturae 104, 199–209. doi:10.1016/j.scienta.2004.08.013.

Newsham, K. K. (2011). A meta-analysis of plant responses to dark septate root endophytes. The New phytologist 190, 783–93. doi:10.1111/j.1469-8137.2010.03611.x.

Nguyen, N. H., Song, Z., Bates, S. T., Branco, S., Tedersoo, L., Menke, J., et al. (2016). FUNGuild: An open annotation tool for parsing fungal community datasets by ecological guild. Fungal Ecology 20, 241–248. doi:10.1016/j.funeco.2015.06.006.

Olivain, C., Humbert, C., Nahalkova, J., Fatehi, J., L’Haridon, F., and Alabouvette, C. (2006). Colonization of tomato root by pathogenic and nonpathogenic Fusarium oxysporum strains inoculated together and separately into the soil. Applied and Environmental Microbiology 72, 1523–1531. doi:10.1128/AEM.72.2.1523-1531.2006.

Paredes, H. S., Gao, T., Law, T. F., Finkel, O. M., Mucyn, T., Teixeira, P. J. P. L., et al. (2018). Design of synthetic bacterial communities for predictable plant phenotypes. PLoS Biology 16, 1–41. doi:10.1371/journal.pbio.2003962.

Peay, K. G., Kennedy, P. G., and Talbot, J. M. (2016). Dimensions of biodiversity in the Earth mycobiome. Nature Reviews Microbiology 14, 434–447. doi:10.1038/nrmicro.2016.59.

Pieterse, C. M. J., Zamioudis, C., Berendsen, R. L., Weller, D. M., van Wees, S. C. M., and Bakker, P. A. H. M. (2014). Induced Systemic Resistance by Beneficial Microbes. Annual Review of Phytopathology 52, 347–375. doi:10.1146/annurev-phyto-082712-102340.

Pineda, A., Dicke, M., Pieterse, C. M. J., and Pozo, M. J. (2013). Beneficial microbes in a changing environment: Are they always helping plants to deal with insects? Functional Ecology 27, 574–586. doi:10.1111/1365-2435.12050.

Priyadharsini, P., and Muthukumar, T. (2017). The root endophytic fungus Curvularia geniculata from Parthenium hysterophorus roots improves plant growth through phosphate solubilization and phytohormone production. Fungal Ecology 27, 69–77. doi:10.1016/j.funeco.2017.02.007.

Radhakrishnan, R., Khan, A. L., Kang, S. M., and Lee, I. J. (2015). A comparative study of phosphate solubilization and the host plant growth promotion ability of Fusarium verticillioides RK01 and Humicola sp. KNU01 under salt stress. Annals of Microbiology 65, 585–593. doi:10.1007/s13213-014-0894-z.

Remy, W., Taylor, T. N., Hass, H., and Kerp, H. (1994). Four hundred-million-year-old vesicular arbuscular mycorrhizae. Proceedings of the National Academy of Sciences of the United States of America 91, 11841–11843. doi:10.1073/pnas.91.25.11841.

Richardson, A. E., Barea, J. M., McNeill, A. M., and Prigent-Combaret, C. (2009). Acquisition of phosphorus and nitrogen in the rhizosphere and plant growth promotion by microorganisms. Plant and Soil 321, 305–339. doi:10.1007/s11104-009-9895-2.

Rudgers, J. A., Afkhami, M. E., Bell-Dereske, L., Chung, Y. A., Crawford, K. M., Kivlin, S. N., et al. (2020). Climate Disruption of Plant-Microbe Interactions. Annual Review of Ecology, Evolution, and Systematics 51, 561–586. doi:10.1146/annurev-ecolsys-011720-090819.

Saad, M. M., Eida, A. A., and Hirt, H. (2020). Tailoring plant-associated microbial inoculants in agriculture: a roadmap for successful application. Journal of Experimental Botany 71, 3878–3901. doi:10.1093/jxb/eraa111.

Schlaeppi, K., and Bulgarelli, D. (2015). The Plant Microbiome at Work. Molecular Plant-Microbe Interactions MPMI 212, 212–217. doi:10.1094/MPMI-10-14-0334-FI.

Taurian, T., Anzuay, M. S., Angelini, J. G., Tonelli, M. L., Ludueña, L., Pena, D., et al. (2010). Phosphate-solubilizing peanut associated bacteria: Screening for plant growth-promoting activities. Plant and Soil 329, 421–431. doi:10.1007/s11104-009-0168-x.

Taylor, T. N., Remy, W., Haas, H., and Kerp, H. (1995). Fossil arbuscular mycorrhizae from the early Devonian. Mycologia 87, 560–573. doi:10.2307/3760776.

Tedersoo, L., May, T. W., and Smith, M. E. (2010). Ectomycorrhizal lifestyle in fungi: Global diversity, distribution, and evolution of phylogenetic lineages. Mycorrhiza 20, 217–263. doi:10.1007/s00572-009-0274-x.

Toju, H., Abe, M. S., Ishii, C., Hori, Y., Fujita, H., and Fukuda, S. (2020). Scoring species for synthetic community design: network analyses of functional core microbiomes. Frontiers in Microbiology 11, 1361–1361. doi:10.3389/fmicb.2020.01361.

Toju, H., Peay, K. G. K. G., Yamamichi, M., Narisawa, K., Hiruma, K., Naito, K., et al. (2018a). Core microbiomes for sustainable agroecosystems. Nature Plants 4, 247–257. doi:10.1038/s41477-018-0139-4.

Toju, H., Tanabe, A. S., and Sato, H. (2018b). Network hubs in root-associated fungal metacommunities. Microbiome 6, 116. doi:10.1101/270371.

Toju, H., Yamamichi M., Jr, P. R. G., Olesen, J. M., Mougi, A., Yoshida, T., et al. (2017). Species-rich networks and eco-evolutionary synthesis at the metacommunity level. Nature Ecology & Evolution 1, 1–11. doi:10.1038/s41559-016-0024.

Toju, H., Yamamoto, S., Tanabe, A. S., Hayakawa, T., and Ishii, H. S. (2016). Network modules and hubs in plant-root fungal biomes. Journal of the Royal Society Interface 13, 20151097–20151097. doi:10.1098/rsif.2015.1097.

Trivedi, P., Leach, J. E., Tringe, S. G., Sa, T., and Singh, B. K. (2020). Plant–microbiome interactions: from community assembly to plant health. Nature Reviews Microbiology 18, 607–621. doi:10.1038/s41579-020-0412-1.

Tsolakidou, M. D., Stringlis, I. A., Fanega-Sleziak, N., Papageorgiou, S., Tsalakou, A., and Pantelides, I. S. (2019). Rhizosphere-enriched microbes as a pool to design synthetic communities for reproducible beneficial outputs. FEMS Microbiology Ecology 95, 1–14. doi:10.1093/femsec/fiz138.

Usuki, F., and Narisawa, K. (2007). A mutualistic symbiosis between a dark septate endophytic fungus, Heteroconium chaetospira, and a nonmycorrhizal plant, Chinese cabbage. Mycologia 99, 175–184. doi:10.3852/mycologia.99.2.175.

van der Heijden, M. G. A., and Hartmann, M. (2016). Networking in the plant microbiome. PLoS Biology 14, 1–9. doi:10.1371/journal.pbio.1002378.

van der Heijden, M. G. A., Klironomos, J. N., Ursic, M., Moutoglis, P., Streitwolf-Engel, R., Boller, T., et al. (1998). Mycorrhizal fungal diversity determines plant biodiversity, ecosystem variability and productivity. Nature 396, 69–72. doi:10.1038/23932.

van Wees, S. C., van der Ent, S., and Pieterse, C. M. (2008). Plant immune responses triggered by beneficial microbes. Current Opinion in Plant Biology 11, 443–448. doi:10.1016/j.pbi.2008.05.005.

Vinale, F., Sivasithamparam, K., Ghisalberti, E. L., Marra, R., Woo, S. L., and Lorito, M. (2008). Trichoderma–plant–pathogen interactions. Soil Biology and Biochemistry 40, 1–10. doi:10.1016/j.soilbio.2007.07.002.

Vorholt, J. A., Vogel, C., Carlström, C. I., and Müller, D. B. (2017). Establishing causality: opportunities of dynthetic vommunities for plant microbiome tesearch. Cell Host and Microbe 22, 142–155. doi:10.1016/j.chom.2017.07.004.

Wagg, C., Schlaeppi, K., Banerjee, S., Kuramae, E. E., and van der Heijden, M. G. A. (2019). Fungal-bacterial diversity and microbiome complexity predict ecosystem functioning. Nature Communications 10, 1–10. doi:10.1038/s41467-019-12798-y.

Wang, X., Pan, Q., Chen, F., Yan, X., and Liao, H. (2011). Effects of co-inoculation with arbuscular mycorrhizal fungi and rhizobia on soybean growth as related to root architecture and availability of N and P. Mycorrhiza 21, 173–181. doi:10.1007/s00572-010-0319-1.

Ważny, R., Rozpądek, P., Jędrzejczyk, R. J., Śliwa, M., Stojakowska, A., Anielska, T., et al. (2018). Does co-inoculation of Lactuca serriola with endophytic and arbuscular mycorrhizal fungi improve plant growth in a polluted environment? Mycorrhiza 28, 235–246. doi:10.1007/s00572-018-0819-y.

Wei, Z., Gu, Y., Friman, V.-P. P., Kowalchuk, G. A., Xu, Y., Shen, Q., et al. (2019). Initial soil microbiome composition and functioning predetermine future plant health. Science Advances 5, 1–11. doi:10.1126/sciadv.aaw0759.

Werner, G. D. A., and Kiers, E. T. (2015). Order of arrival structures arbuscular mycorrhizal colonization of plants. New Phytologist 205, 1515–1524. doi:10.1111/nph.13092.

Zhu, J., Tremblay, N., and Liang, Y. (2012). Comparing SPAD and atLEAF values for chlorophyll assessment in crop species. Canadian Journal of Soil Science 92, 645–648. doi:10.4141/CJSS2011-100.

