## Supplementary Fig. for "Synergistic and offset effects of fungal species combinations on plant performance"

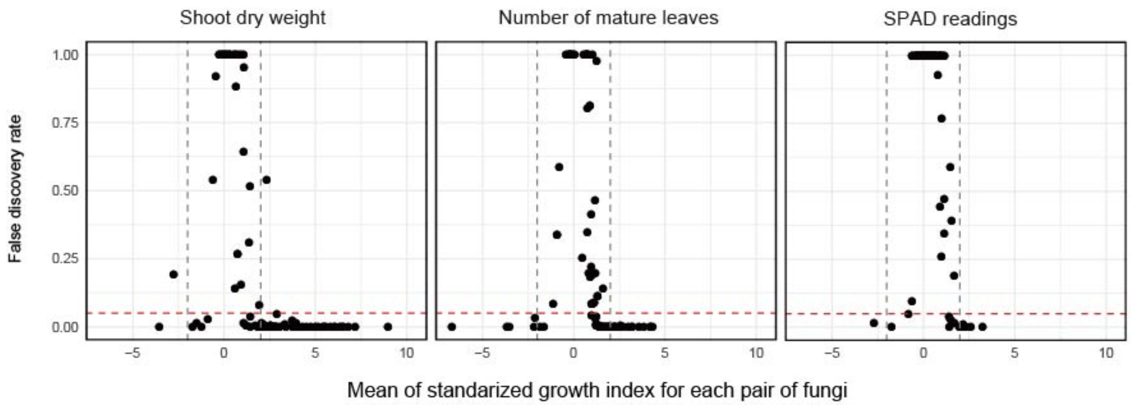


**Supplementary Figure S1 |** Standardized growth index and its statistical sign. The false discovery rates of *t*-tests between control and target single-/dual-inoculation treatments are plotted against the axis of standardized growth index values averaged across replicate samples.

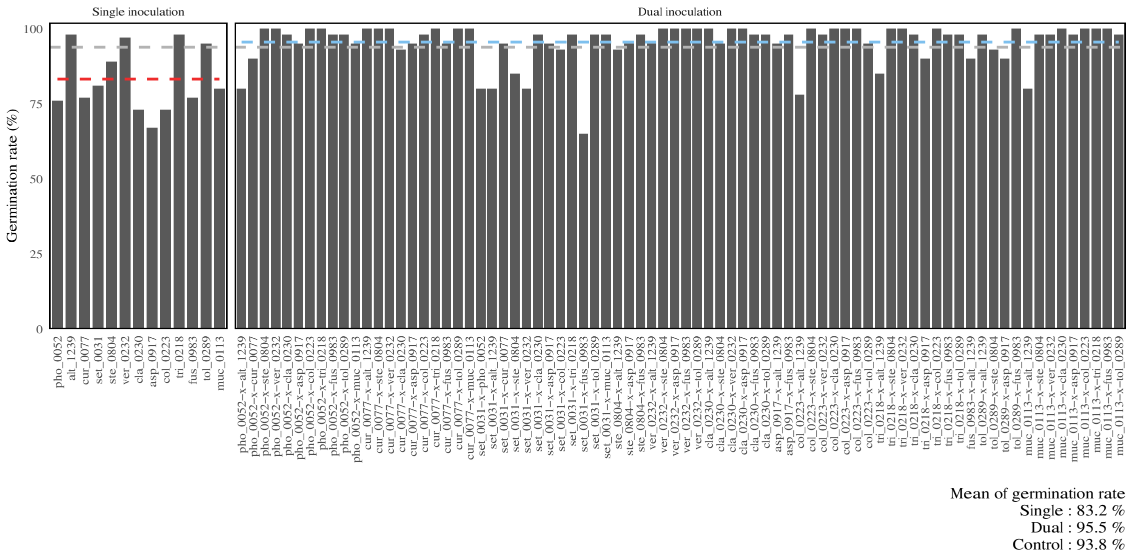


**Supplementary Figure S2 |** Germination rates of *Brassica* plants in the single- and dual-inoculation experiments.

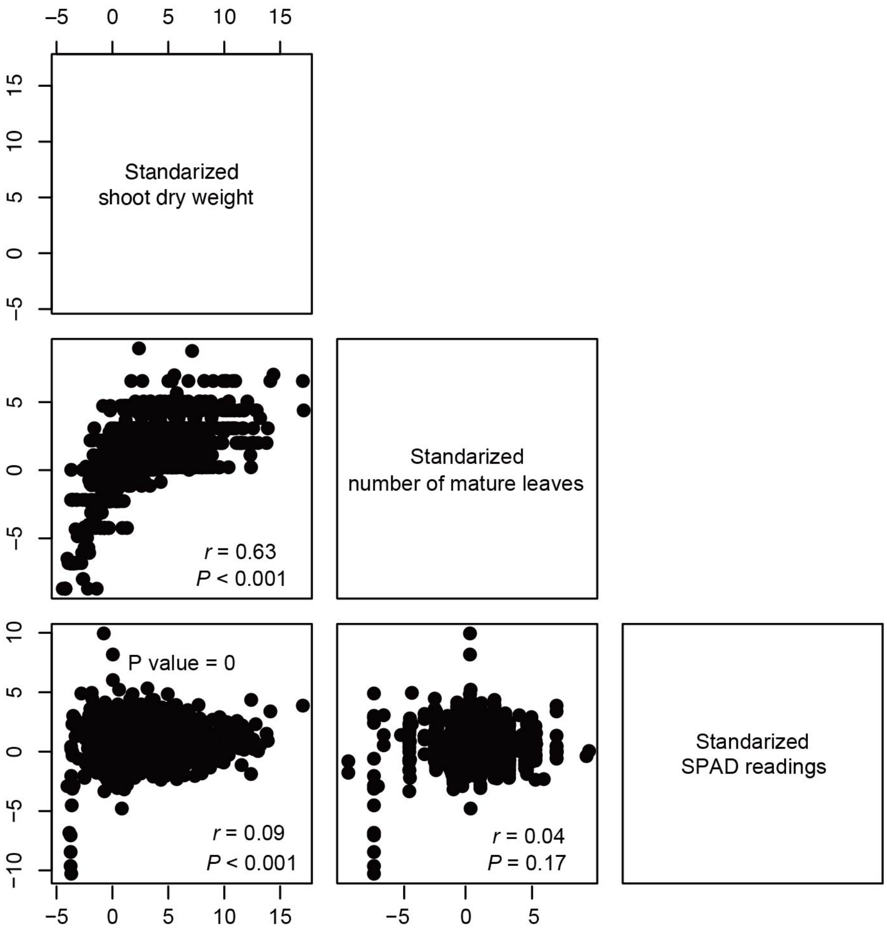


**Supplementary Figure S3 |** Relationships among plant performance traits.
